# CCCP experimental evolution of *Escherichia coli* selects for mutations that increase EmrA activity or that downregulate other PMF-driven drug efflux pumps

**DOI:** 10.1101/392977

**Authors:** Jessie M. Griffith, Preston J. Basting, Katarina M. Bischof, Erintrude P. Wrona, Karina S. Kunka, Anna C. Tancredi, Jeremy P. Moore, Miriam R. L. Hyman, Joan L. Slonczewski

**Affiliations:** Department of Biology, Kenyon College, Gambier, Ohio, USA

## Abstract

**ABSTRACT:** Experimental evolution was conducted with *Escherichia coli* K-12 W3110 in the presence of carbonyl cyanide m-chlorophenylhydrazone (CCCP), an uncoupler of the proton motive force (PMF). Cultures were serially diluted daily 1:100 in broth medium containing 20-150 μM CCCP at pH 6.5 or at pH 8.0. After 1,000 generations, all populations showed 5- to 10-fold increase in CCCP resistance. Sequenced isolates showed mutations in *emrAB* or in its negative repressor *mprA*; the EmrAB-TolC multidrug efflux pump confers resistance to CCCP and nalidixic acid. Deletion of *emrA* abolished the CCCP resistance of these strains. One CCCP-evolved isolate lacked *emrA* or *mprA* mutations; this strain (C-B11-1) showed mutations in drug efflux regulators *cecR* (*ybiH*) (upregulates drug pumps YbhG and YbhFSR) and *gadE* (upregulates drug pump *mdtEF*). A *cecR∷kanR* deletion conferred partial resistance to CCCP. A later evolved descendant of the C-B11 population showed mutations in *ybhR* (MDR efflux). Another isolate showed *acrB* (MDR efflux pump). The *acrB* isolate was sensitive to chloramphenicol and tetracycline, which are effluxed by AcrAB. Other mutant genes in CCCP-evolved strains include *adhE* (alcohol dehydrogenase), *rng* (ribonuclease G), and *cyaA* (adenylate cyclase). Overall, experimental evolution revealed a CCCP fitness advantage for mutations increasing its own efflux via EmrA; and for mutations that may decrease proton-driven pumps that efflux other drugs not present (*cecR, gadE, acrB, ybhR*). These results are consistent with our previous report of drug sensitivity associated with evolved tolerance to a partial uncoupler (benzoate or salicylate).

**IMPORTANCE:** The genetic responses of bacteria to depletion of proton motive force, and their effects on drug resistance, are poorly understood. Our evolution experiment reveals genetic mechanisms of adaptation to the PMF uncoupler CCCP, including selection for and against various multidrug efflux pumps. The results have implications for our understanding of the gut microbiome, which experiences high levels of organic acids that decrease PMF. Organic acid uncouplers may select against multidrug resistance in evolving populations of enteric bacteria.

## INTRODUCTION

The proton motive force (PMF) is diminished or abolished by uncouplers of oxidative phosphorylation such as carbonyl cyanide m-chlorophenyl hydrazone (CCCP) (1). Uncouplers such as CCCP are generally hydrophobic compounds with an acidic proton that reside in the inner membrane of the cell. These cations then shuttle protons across the membrane via protonation/deprotonation (1–3). The transmembrane proton flux equilibrates both ΔpH and Δψ, and thus depletes PMF (1, 2). Respiration then runs a futile cycle, as protons continue to be pumped but PMF is not maintained (4–6). The uncoupler effect is most pronounced during growth at low external pH (pH 5.0-5.7) where the electron transport system is upregulated and a higher PMF is maintained (7).

Evolution experiments show how *E. coli* evolve over generations of pH-related stress and increase fitness under low pH, high pH, and benzoic acid exposure (8–11). Such experiments often reveal surprising fitness tradeoffs. Evolution under continual low pH exposure leads to loss of activity of amino acid decarboxylases that are highly induced under short-term acid stress (8, 10). Evolution in benzoic acid, a membrane permeant aromatic acid, yields unexpected loss of multidrug resistance (MDR) efflux pumps and antibiotic resistance (9). For example, benzoate stress selects for mutations that delete or downregulate the Gad acid fitness island including the *mdtEF* drug efflux system (9, 12). At high concentration, benzoic acid partly uncouples PMF and thus could increase the fitness cost of efflux pumps driven by proton flux.

It was of interest therefore to test the fitness effect of long-term exposure to a strong uncoupler, CCCP, that more completely abolishes PMF. One system of interest for CCCP tolerance is EmrAB-TolC. EmrA, EmrB, and TolC form a multidrug efflux pump that exports CCCP and various ionophores and antibiotics (13–15). The *emrAB* operon is regulated by MprA, a repressor of the operon under the same promoter control as *emrAB* (16). MprA binds CCCP and becomes inactivated, allowing for higher expression and activity of EmrA and EmrB. It is unknown whether long-term CCCP exposure would select for increased activity of the pump, or its regulators; or for loss of this CCCP-responsive system, as is found for the loss of acid-inducible decarboxylases following long-term exposure to acid (10).

We performed experimental evolution to test the long-term effects of exposure to a full uncoupler, and the role of external pH in CCCP tolerance. We conducted serial dilution of *E. coli* at pH 6.5 and 8.0 with increasing concentrations of CCCP.

## RESULTS

### CCCP-evolved populations show increased relative fitness in the presence of CCCP

To investigate the selection effects of CCCP on *E. coli,* populations of strain W3110 were subcultured daily with CCCP in medium buffered at pH 6.5 or at pH 8.0 (**Table 1**). The initial CCCP concentrations, 20 μM for low pH and 50 μM for high pH, respectively, were determined by culturing W3110 in a range of CCCP concentrations at pH 6.5 and pH 8.0 (**Figs. 1A and 1B**). The ancestral strain W3110 failed to grow consistently above 40 μM CCCP (pH 6.5) or above 60 μM CCCP (pH 8.0). Culture densities at 16 h showed that 20 μM CCCP at pH 6.5 and 50 μM CCCP at pH 8.0 resulted in significant decrease of growth without full loss of viability. From this point forward, CCCP concentration was increased in a stepwise fashion, reaching 150 μM CCCP by generation 1,000. **Figure 1C** compares the 16-h endpoint culture densities attained by the evolving populations versus the ancestral strain W3110, using samples from populations frozen at succeeding generations up to 1,000. The pH 8.0 populations showed a steeper increase in fitness than those exposed to CCCP at pH 6.5, where fitness leveled off after 600 generations.

**Figure 1.**
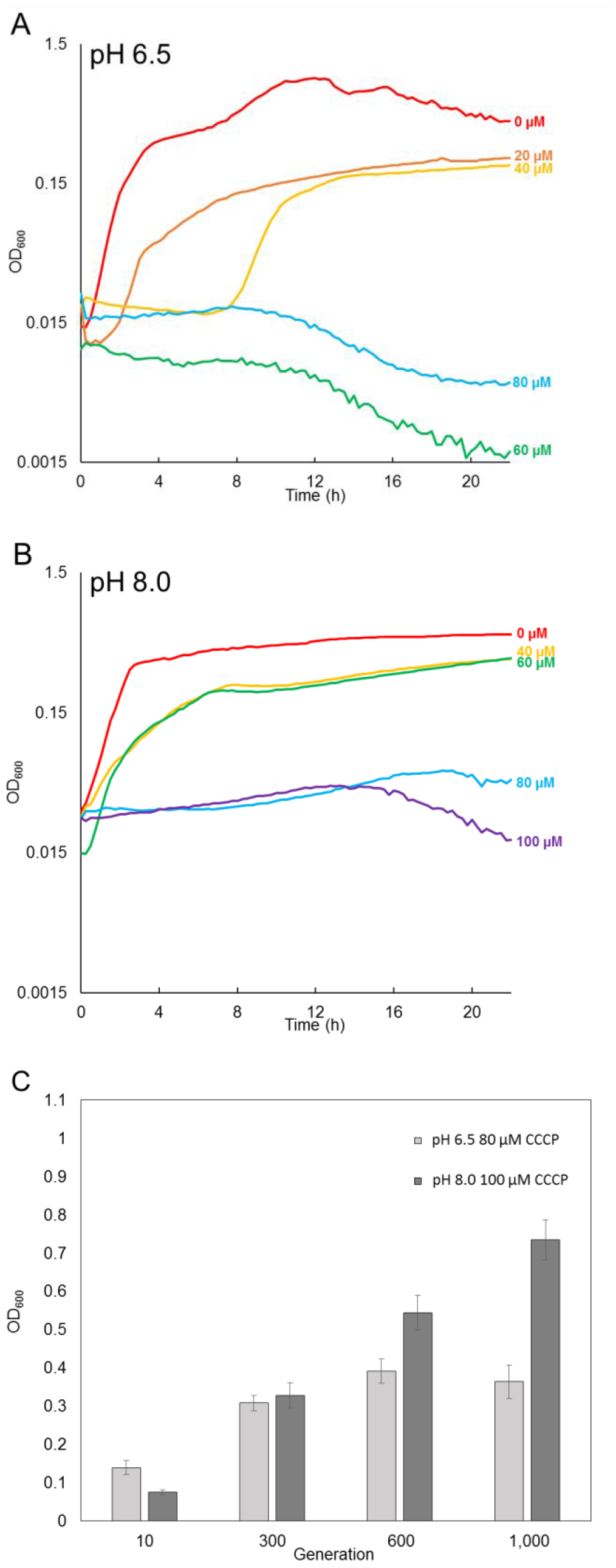
Culture densities attained by populations of strain W3110 serially diluted and exposed to increasing CCCP concentrations. Microplate 200-μl populations were diluted in media containing (**A**) 100 mM Na-PIPES pH 6.5 and (**B**) 100 mM TAPS pH 8.0. (**C**) Mean endpoints at t=16 h are shown for evolving populations cultured with 80 μM CCCP at pH 6.5 (light bars), and in 100 μM CCCP at pH 8.0 (dark bars). At pH 6.5, 80 μM CCCP, 8 samples were excluded from the generation 10 OD600 data because no growth occurred.

**Table 1.** Generation number at increases in CCCP concentration during evolution of *E. coli*

| Generation number: | New concentration CCCP (μM) for pH 6.5 condition | New concentration CCCP (μM) for pH 8.0 condition |
| --- | --- | --- |
| 0 | 20 | 60 |
| 20 | 20 | 50 |
| 66 | 30 | 60 |
| 299 | 40 | 70 |
| 345 | 50 | 80 |
| 523 | 60 | 90 |
| 536 | 70 | 100 |
| 556 | 80 | 110 |
| 616 | 120 | 130 |
| 782 | 150 | 150 |

After 1,000 generations, isolates were obtained from selected microplate populations. Isolated strains are named by the position on the plate and isolate number; for example, strain C-A1-1 was the first CCCP-evolved strain from the well in row A and column 1. Strain names for each strain are listed in **Table 2**. **Figure 2** shows growth curves obtained for isolates from populations following evolution at pH 8.0 (panels A, B) or at pH 6.5 (panels C, D). For each isolate, eight replicate curves were obtained. Panels A and C show the curve exhibiting median density at 16 h for each strain and condition; panels B and D show all eight replicate curves. Isolates that had evolved at pH 8.0 (C-B11-1, C-D11-1, C-F9-1, C-G7-1) as well as isolates that had evolved at pH 6.5 (C-A1-1, C-DA3-1, C-G5-1) showed an increase in tolerance to 150 μM CCCP.

**Figure 2.**
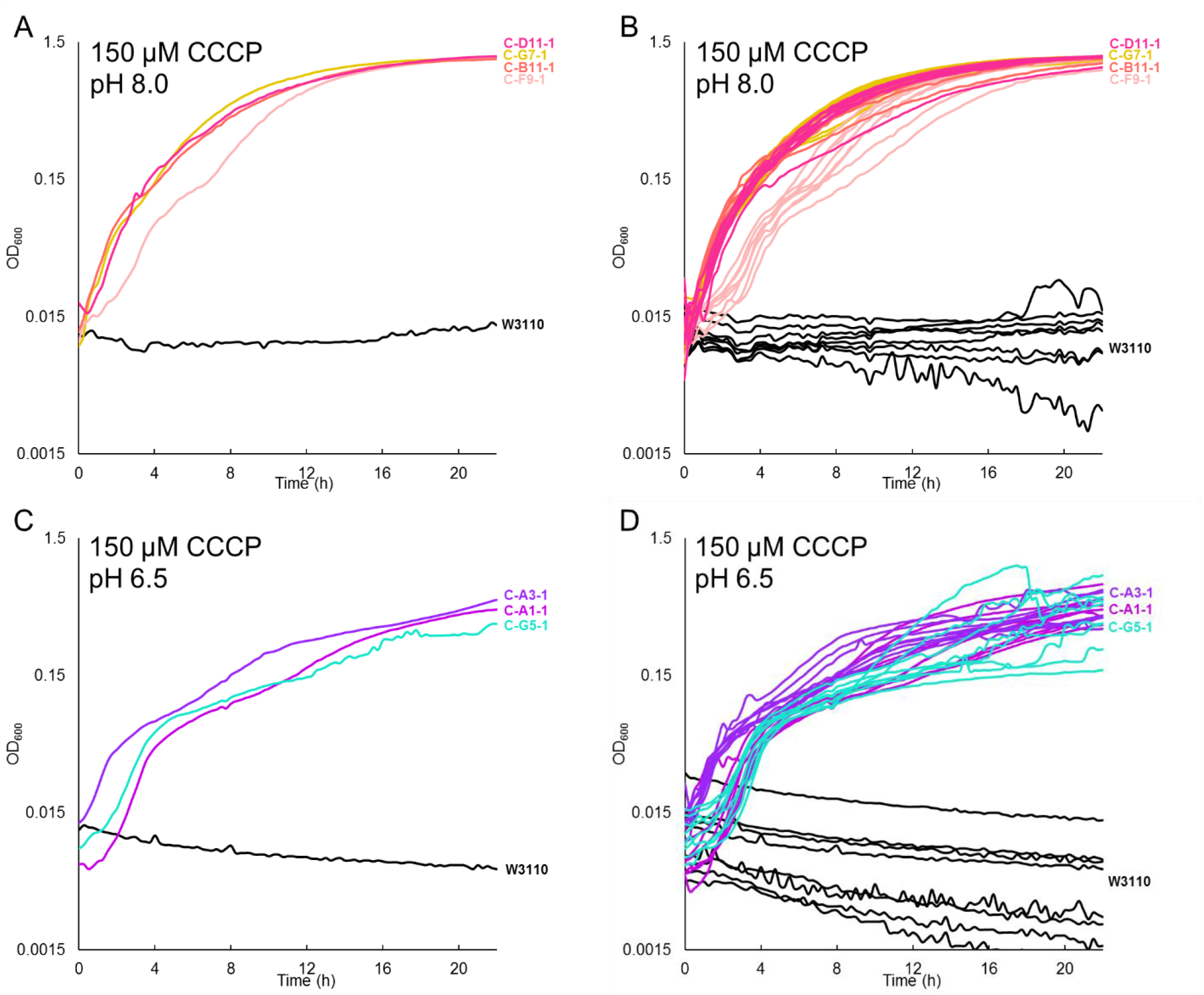
Isolates from CCCP-evolved populations show increased tolerance for 150 μM CCCP. (**A and B**) medium buffered with 100 mM TAPS, pH 8.0; (**C and D**) 100 mM Na-PIPES, pH 6.5. Panels A and C show representative curves with median density at 16 h; panels B and D show all 8 replicates.

**Table 2.** Strains generated by experimental evolution or by P1 phage transduction

| Strain | Population isolate | Description or Genotype | Generation |
| --- | --- | --- | --- |
| W3110 | -- | *Escherichia coli* K-12 | -- |
| JLSC0001 | C-A1-1 | W3110 evolved at pH 6.5 with CCCP | 1,000 |
| JLSC0005 | C-E1-1 | W3110 evolved at pH 6.5 with CCCP | 1,000 |
| JLSC0009 | C-A3-1 | W3110 evolved at pH 6.5 with CCCP | 1,000 |
| JLSC0010 | C-B3-1 | W3110 evolved at pH 6.5 with CCCP | 1,000 |
| JLSC0013 | C-E3-1 | W3110 evolved at pH 6.5 with CCCP | 1,000 |
| JLSC0016 | C-H3-1 | W3110 evolved at pH 6.5 with CCCP | 1,000 |
| JLSC0017 | C-A5-1 | W3110 evolved at pH 6.5 with CCCP | 1,000 |
| JLSC0023 | C-G5-1 | W3110 evolved at pH 6.5 with CCCP | 1,000 |
| JLSC0024 | C-H5-1 | W3110 evolved at pH 6.5 with CCCP | 1,000 |
| JLSC0028 | C-D7-1 | W3110 evolved at pH 8.0 with CCCP | 1,000 |
| JLSC0031 | C-G7-1 | W3110 evolved at pH 8.0 with CCCP | 1,000 |
| JLSC0033 | C-A9-1 | W3110 evolved at pH 8.0 with CCCP | 1,000 |
| JLSC0038 | C-F9-1 | W3110 evolved at pH 8.0 with CCCP | 1,000 |
| JLSC0042 | C-B11-1 | W3110 evolved at pH 8.0 with CCCP | 1,000 |
| JLSC0044 | C-D11-1 | W3110 evolved at pH 8.0 with CCCP | 1,000 |
| JLSC0058 | C-B11-3 | W3110 evolved at pH 8.0 with CCCP | 2,000 |
| JLSC0059 | C-B11-4 | W3110 evolved at pH 8.0 with CCCP | 2,000 |
| JLS1718 | -- | W3110 $\Delta mprA::kanR$ | -- |
| JLS1740 | -- | W3110 $\Delta ybiH::kanR$ | -- |
| JLS1809 | -- | W3110 $\Delta adhE::kanR$ | -- |
| JLS1706 | -- | W3110 $\Delta emrA::kanR$ | -- |
| JLS1701 | -- | JLSC0001 (C-A1-1) $\Delta emrA::kanR$ | -- |
| JLS1702 | -- | JLSC0009 (C-A3-1) $\Delta emrA::kanR$ | -- |
| JLS1703 | -- | JLSC0023 (C-G5-1) $\Delta emrA::kanR$ | -- |
| JLS1704 | -- | JLSC0042 (C-B11-1) $\Delta emrA::kanR$ | -- |
| JLS1705 | -- | JLSC0044 (C-D11-1) $\Delta emrA::kanR$ | -- |
| JLS1733 | -- | JLSC0042 (C-B11-1) *rng*<sup>+</sup> | -- |

### Genomes of CCCP-evolved strains show independent recurring mutations in common genes

The genomes of CCCP-evolved strains were resequenced and analyzed using the computational pipeline *breseq* to characterize mutations acquired over the course of the evolution (**Table 3**, selected mutations; **Table S1**, all mutations). Isolate C-B3-1 behaved as a mutator, showing approximately 20-fold higher mutation rate than the other strains; this strain contained a mutation to *mutS* (**Table S2**). Isolates C-E1-1 and C-A1-1 were genetically highly similar, indicating that the strains were nearly clones. For these reasons, isolates C-B3-1 and C-E1-1 were excluded from further study.

For eight CCCP-evolved populations at pH 6.5, and five at pH 8.0, all mutations predicted by *breseq* are shown in **Table S1**. Selected mutations, including all for genes that showed mutations in more than one population and for genes of the *emrAB* operon or its repressor, are shown in **Table 3**. The table is organized by condition, with mutations found in populations cultured at pH 6.5 are shaded blue, whereas mutations found at high pH are shaded red. Mutations that occurred in both the low pH and high pH project are shaded purple.

**Table 3.**
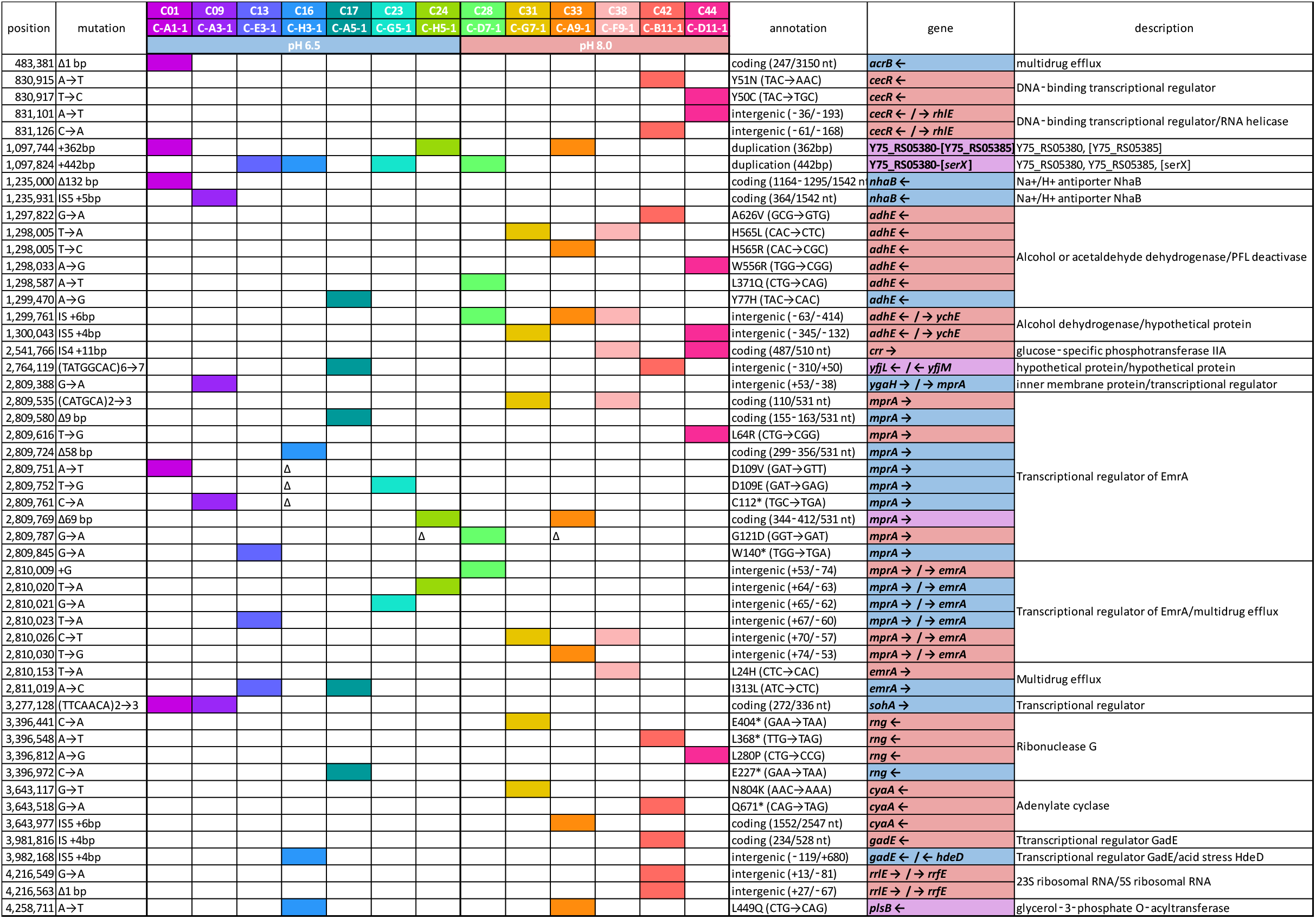
Selected mutations after evolution with CCCP at pH 6.5 or at pH 8.0.^a^

All populations that evolved with CCCP at pH 6.5, and five out of six that evolved at pH 8.0, showed mutations in *mprA,* which encodes the repressor of *emrAB* (**Table 3**) (13, 17). Of the 12 strains with mutations in *mprA,* there were a total of 10 unique mutations to the coding region of MprA and 6 mutations to the intergenic region between *mprA* and *emrA.* Two mutations caused early stop codons in MprA, and two other mutations caused deletions of 50 bp or greater (**Table 3**). MprA represses the *emrAB* operon which encodes a multidrug efflux pump which exports CCCP from the cell (14, 15, 18). Thus, *mprA* repressor knockouts could be associated with increased production of the EmrAB ad TolC multidrug efflux pump, and could mediate the increased CCCP tolerance in the CCCP-evolved strains. Nine of the thirteen strains show additional mutations in *emrAB* or in the intergenic region between *mprA* and *emrA*, which includes promoter control sequences. No knockout insertions or deletions were seen for *emrAB.*

### Deletion of ***mprA*** enhances growth in CCCP

Given the fitness selection for mutations in *mprA,* we tested the role of MprA in CCCP tolerance by deleting *mprA* from the ancestral strain *E. coli* W3110. Both at pH 6.5 and at pH 8.0, the Δ*mprA∷kanR* deletant grew at higher concentration of CCCP than the ancestral strain W3110 (**Fig. 3**). In addition, loss of MprA causes no growth difference in the absence of CCCP. Thus, the growth improvements due to Δ*mprA∷kanR* are associated with CCCP, probably by efflux via EmrAB complex. The increased CCCP tolerance of Δ*mprA∷kanR* is consistent with the hypothesis that loss of MprA repressor function in CCCP-evolved isolates increases fitness in the presence of CCCP.

**Figure 3.**
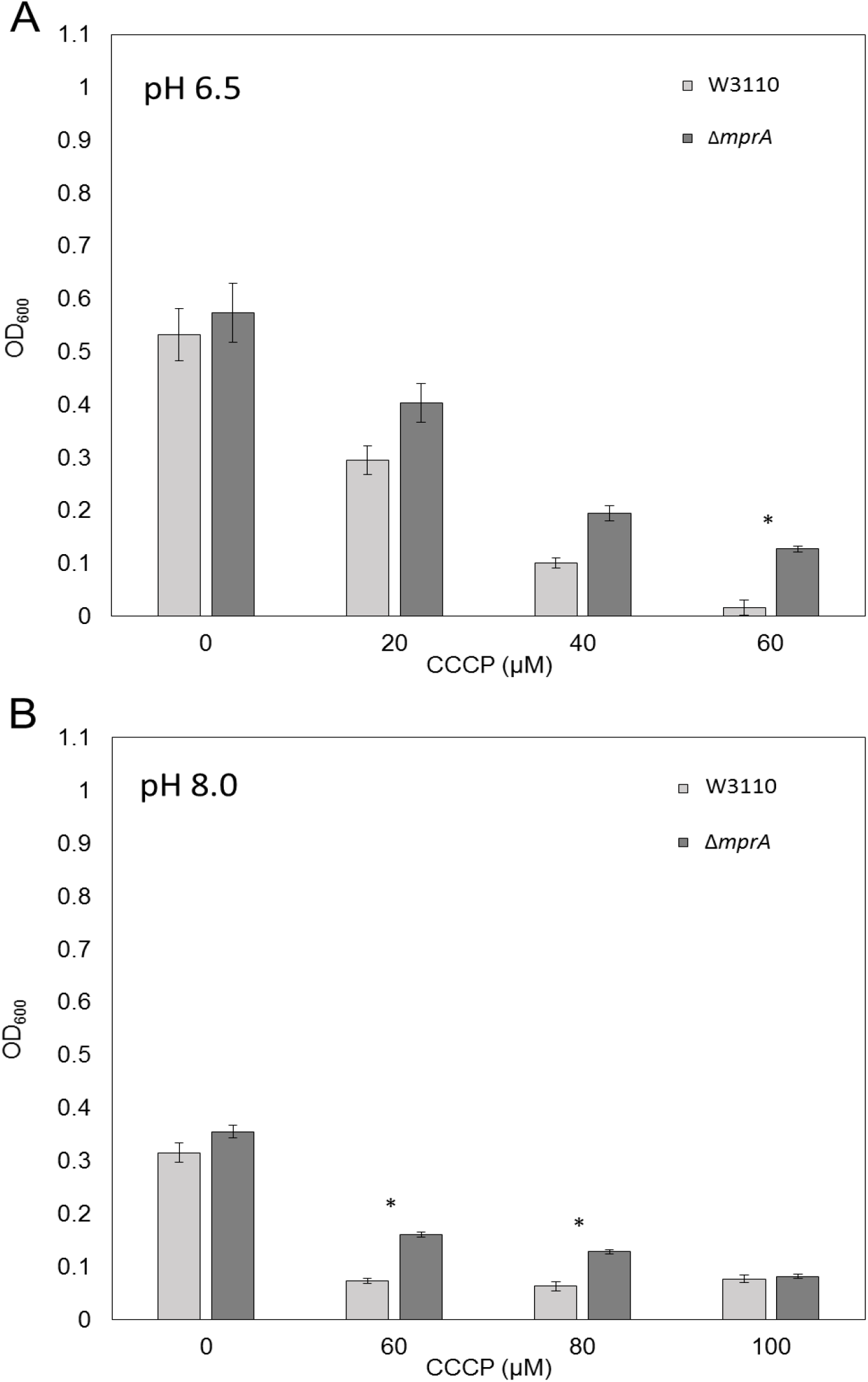
*AmprA* increases resistance of W3110 to CCCP at both low and high pH. W3110Δ*mprA∷kanR* (dark grey) and W3110 (light grey) in increasing concentrations of CCCP at (**A**) LBK 100 mM Na-PIPES pH 6.5 and (**B**) LBK 100 mM TAPS pH 8.0. Asterisk indicates significant Tukey”s test p<0.05.

### Deletion of ***emrA*** in CCCP-evolved strains decreases CCCP fitness

Three of the CCCP-evolved strains (C-E3-1, C-A5-1, C-F9-1) showed missense mutations in *emrA.* We therefore tested whether *emrA* activity was required for these and other CCCP-evolved strains. Deletion of *emrA* (by transduction of *ΔemrA∷kanR*) in strains C-A1-1, C-A3-1, C-G5-1, and C-D11-1 substantially decreased growth compared to the strains with *emrA* intact, both at pH 6.5 and at pH 8.0 with CCCP (**Fig. 4**). Thus, *emrA* appears to be required for most if not all of the fitness advantage of these evolved strains.

**Figure 4.**
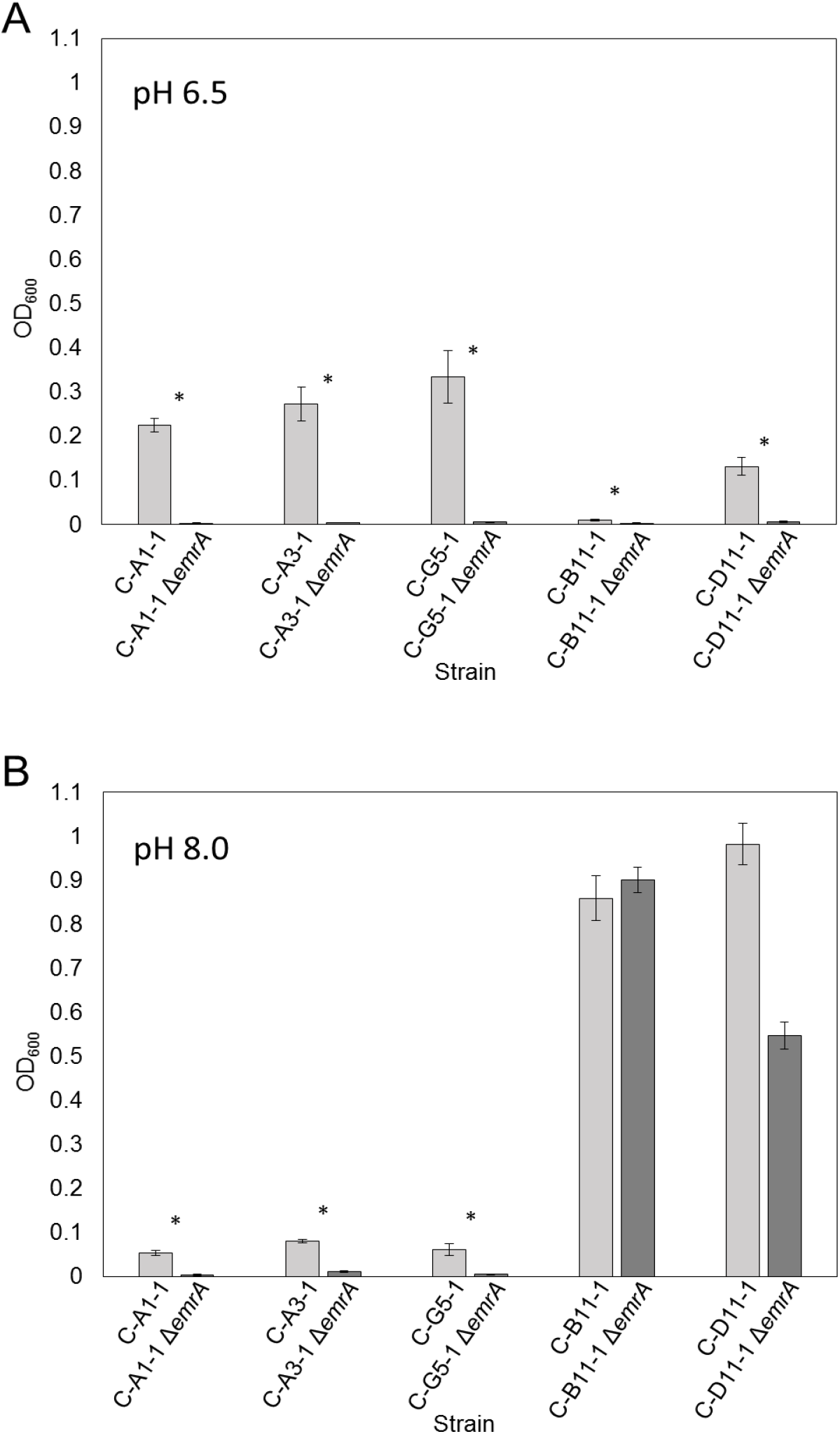
*ΔemrA* decreases growth of CCCP-evolved strains in the presence of 150 μM CCCP. Growth of A1-1, A3-1, G5-1, B11-1, and D11-1 (dark gray) in 150 μM CCCP (**A**) LBK pH 6.5 100 mM Na-PIPES and (**B**) LBK pH 8.0 100 mM TAPS compared to growth of A1-*1ΔemrA∷kanR,* A3-1 *ΔemrA∷kanR,* G5-1 *ΔemrA∷kanR, B11-1ΔemrA∷kanR,* and D11-*1ΔemrA∷kanR* (light grey). Asterisk indicates significant Tukey”s test p<0.05.

The one CCCP-evolved strain that lacked any *emrA* or *mprA* mutation (strain C-B11-1) showed a fitness advantage that was independent of *emrA,* during culture at pH 8.0 (**Fig. 4B**). Strain C-D11-1 *ΔemrA∷kanR* also showed partial CCCP tolerance. However, all strains including C-B11-1 required *emrA* for growth at pH 6.5, where the uncoupler effect of CCCP is greatest.

### Mutations affecting ***cecR*** in strain C-B11-1 increase CCCP tolerance

The CCCP-evolved isolate C-B11-1 contains no mutations in *mprA* or *emrAB,* so we sought to identify other major components of CCCP tolerance in this strain. C-B11-1 notably sustained an insH-mediated deletion of nearly 20 kb which includes *phoE, proA* and *proB, perR, pepD, gpt, frsA,* and *crl* (**Table S1**). In addition to this large deletion, C-B11-1 also showed a mutation to *cecR (ybiH)* and a mutation in the intergenic region between *cecR* and *rhlE*. Mutations to this region were also present in another high pH CCCP-evolved strain, C-D11-1. CecR regulates genes that affect sensitivity to cefoperazone and chloramphenicol (19). We found that W3110 Δ*cecR∷kanR* showed increased CCCP tolerance at pH 6.5 (Fig. 5). In the presence of 30 μM CCCP, the *cecR* deletion strain reached a cell density comparable to that of isolate C-B11-1, whereas the ancestral strain W3110 grew significantly less. Thus, a *cecR* defect could contribute the major part of the CCCP fitness advantage for C-B11-1.

**Figure 5.**
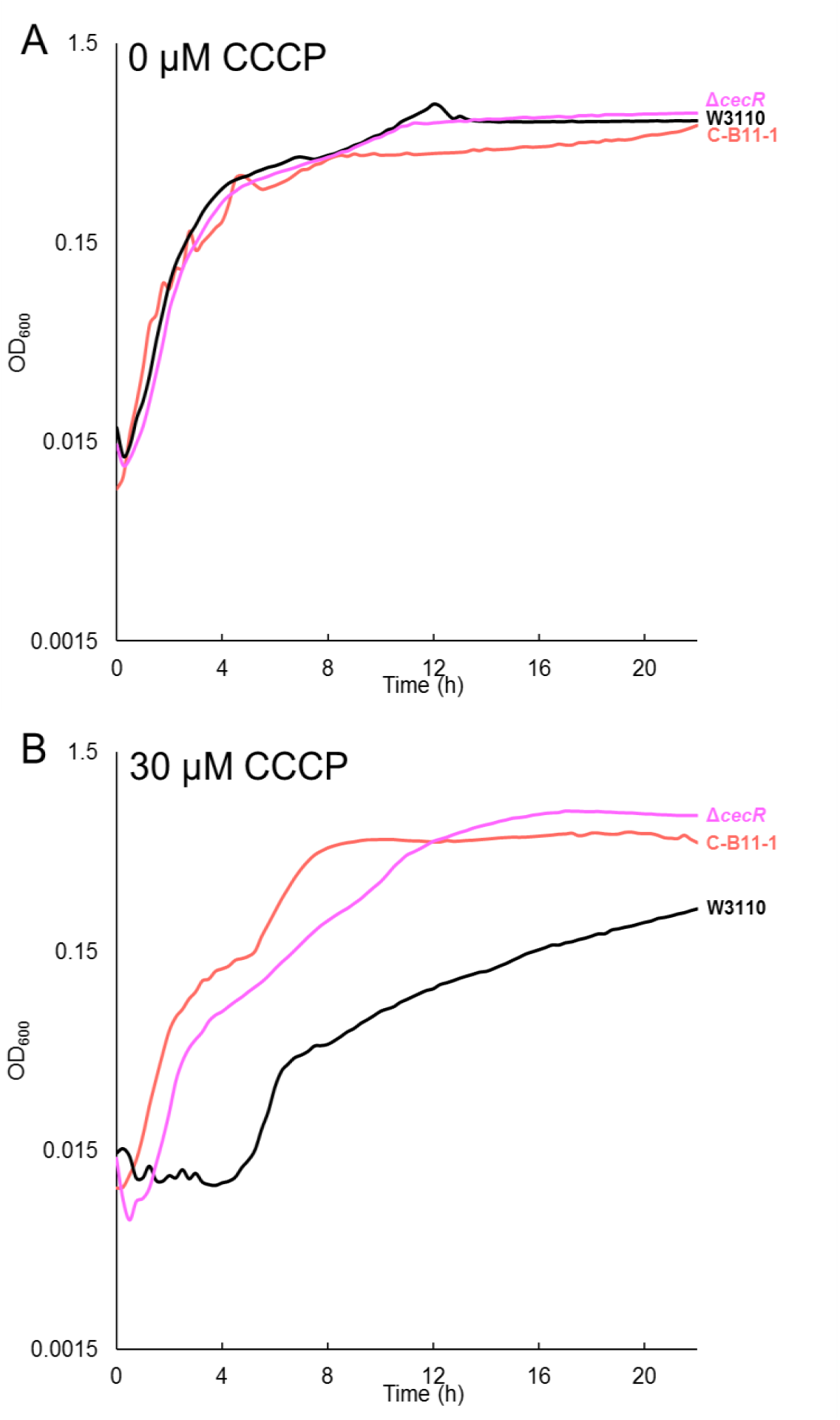
ΔcecR improves growth of W3110 in CCCP stressed condition. Growth of B11-1, *W3110ΔcecR∷kanR,* and W3110 in 100 mM Na-PIPES pH 6.5 (**A**) 0 μM CCCP and (**B**) 30 μM CCCP. (ANOVA with Tukey for (**A**) 0 μM CCCP, F=0.135 and p=0.875. For (**B**) 30 μM CCCP, F=93.63 and p=3.45e^-11^.)

Several additional genes that showed mutations in strain C-B11-1 were tested for effects on CCCP tolerance. A C-B11-1 rng+ construct made by recombineering showed no difference in CCCP tolerance compared to the parental strain C-B11-1, cultured in 150 μM CCCP pH 8.0. Other deletions were tested in the ancestral strain background: W3110 Δ*acrB,* (30 μM CCCP pH 6.5), W3110 Δ*cyaA* (50 μM CCCP pH 6.5) and W3110 Δ*nhaB* (0-50 μM CCCP pH 7.0). None of these constructs showed significant difference in growth compared to strain W3110.

### Additional mutations in population C-B11

The strain C-B11-1 showed additional mutations of interest for drug efflux, most notably *gadE* (20, 21). GadE activates expression of the Gad acid fitness island, which includes drug efflux *mdtFE* (22). The MdtFE efflux pump is not known to transport CCCP. Negative mutation of *gadE* would decrease expression of *mdtFE,* an effect seen in evolution under benzoate stress (9).

We pursued the effect of the C-B11-1 genotype by sequencing two later isolates from the C-B11 population (C-B11-3, C-B11-4) after a total of 2,000 generations (**Table S3**). Cultures of the later isolates showed growth curves comparable to those for C-B11-1. These later isolates retained most of the mutations found in C-B11-1, including the nearly 20-kb deletion of 28 genes that was unique to population C-B11. Between 1,000 and 2,000 generations, the C-B11 strains acquired additional mutations affecting genes *ybhR, lptD, tsx, gltA, cfa, mgrB, pykA, cpsB, ydhN* and *yjjU.* Gene *ybhR* encodes a subunit of the cefoperazone efflux complex regulated by CecR (19). This result is thus consistent with the enhanced fitness shown by *cecR* mutation found in the C-B11 lineage. The *mgrB* gene is acid-inducible, via PhoPQ response (23) and shows mutations selected after high-pH evolution (11). The gene *lptD* forms part of the lipopolysaccharide transport slide (24), which could be relevant for CCCP membrane interactions.

### CCCP-evolved strains show altered resistance to antibiotics

Exposure to the partial uncoupler benzoate selects for strains sensitive to chloramphenicol and tetracycline (9). We therefore tested the growth of the CCCP-evolved strains in the presence of various antibiotics (**Figure 6**). Nalidixic acid is target of the EmrAB TolC multidrug efflux pump (14), so mutations increasing expression of function of this pump might increase resistance to the substrate antibiotics. In the presence of nalidixic acid, several CCCP-evolved strains grew to higher levels than the ancestor (**Figure 6A**). Strain C-B11-1 does not contain mutations known to affect EmrAB, but it contains a mutation affecting CecR, which may increase resistance to nalidixic acid (19).

**Figure 6.**
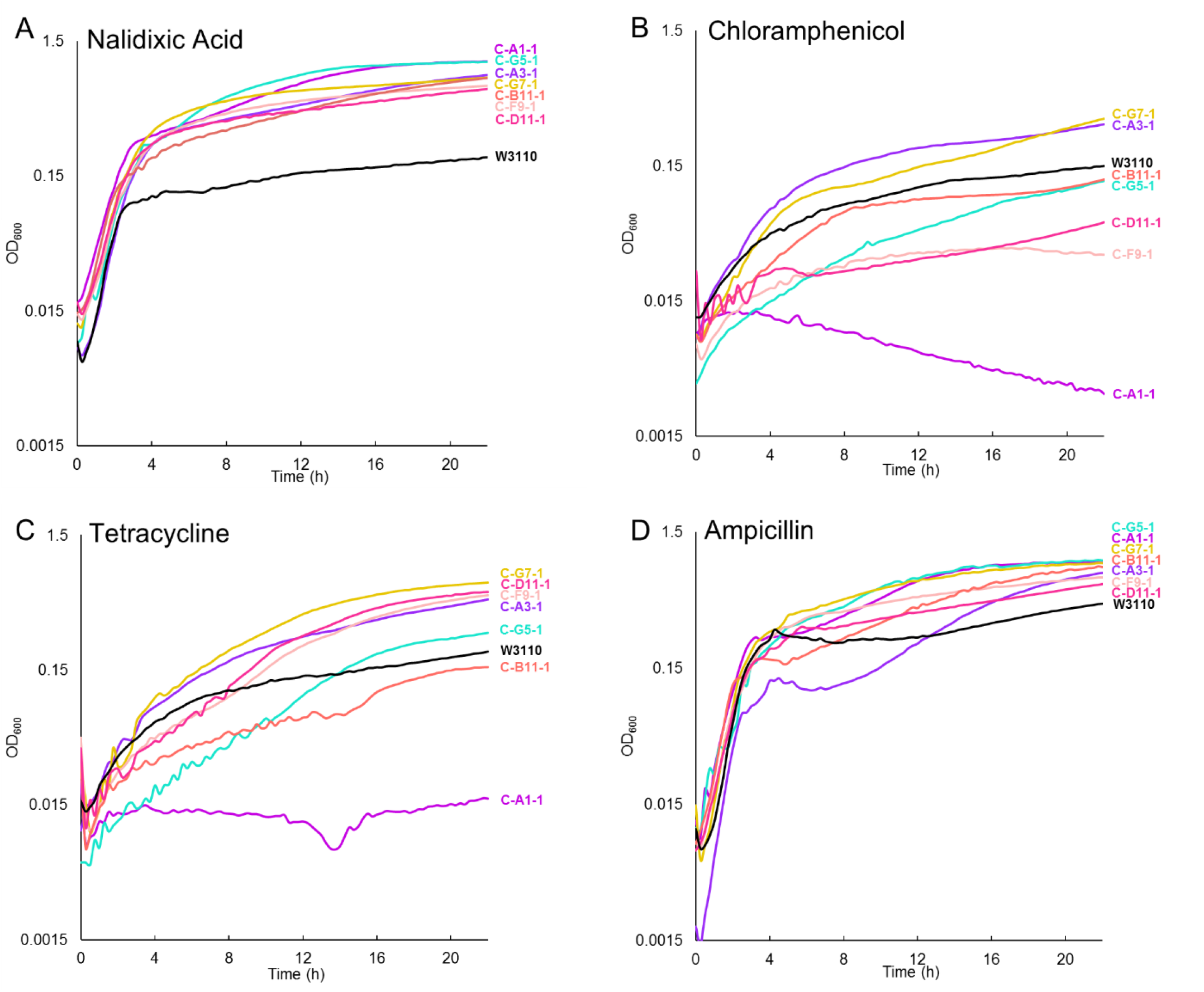
Growth of CCCP-evolved strains compared to the ancestor in LBK 100 mM MOPS pH 7.0 in (**A**) 6 μg/mL nalidixic acid, (**B**) 4 μg/ml chloramphenicol, (**C**) 1 μg/ml tetracycline, and (**D**) 1 μg/ml ampicillin. Curves shown are representative of the 8 replicates for each strain. Endpoint optical density at 16 hours (OD_600_) data as a measure of culture growth compared. For (**A**) growth of all evolved isolates compared to W3110, Tukey gave p < 0.001. For growth of strain C-A1-1 versus W3110 (**B**) in chloramphenicol, and (**C**) in tetracycline, Tukey gave p < 0.001.

Other antibiotics tested for resistance include chloramphenicol (4 μg/ml) (**Figure 6B**) tetracycline (1 μg/ml) (**Figure 6C**), ampicillin (1 μg/ml) (**Figure 6D**). Chloramphenicol, tetracycline and ampicillin showed varied effects in different CCCP-evolved isolates, including marginal gain or loss of resistance. Strain C-A1-1 was very sensitive to chloramphenicol and tetracycline, similar to the result seen for benzoate-evolved strains (9). This strain contained mutations in *acrB* (25, 26), a drug pump conferring resistance to chloramphenicol and tetracycline; also mutations in *sohA* and in *rpoB* that are similar to mutations selected in the benzoate-evolved strains.

### Evolved strains show small pH growth effects independent of CCCP

Serial culture at low or high pH is known to shift the growth range (8, 11). Thus, it was important to test the effect of pH alone during CCCP evolution at low or high pH. The CCCP-evolved strains showed marginal changes in pH dependence, as represented by culture density at 16 h. Strains evolved at pH 6.5 showed growth indistinguishable from that of the ancestral W3110 at pH 6.5 (**Fig. 7A**). At pH 8.0, we saw a small significant increase in density during stationary phase (**Fig. 7B**). Strains evolved at pH 8.0 consistently showed a small increase in stationary phase growth at either pH 6.5 (**Fig. 7C**) or at pH 8.0 (**Fig. 7D**).

**Figure 7.**
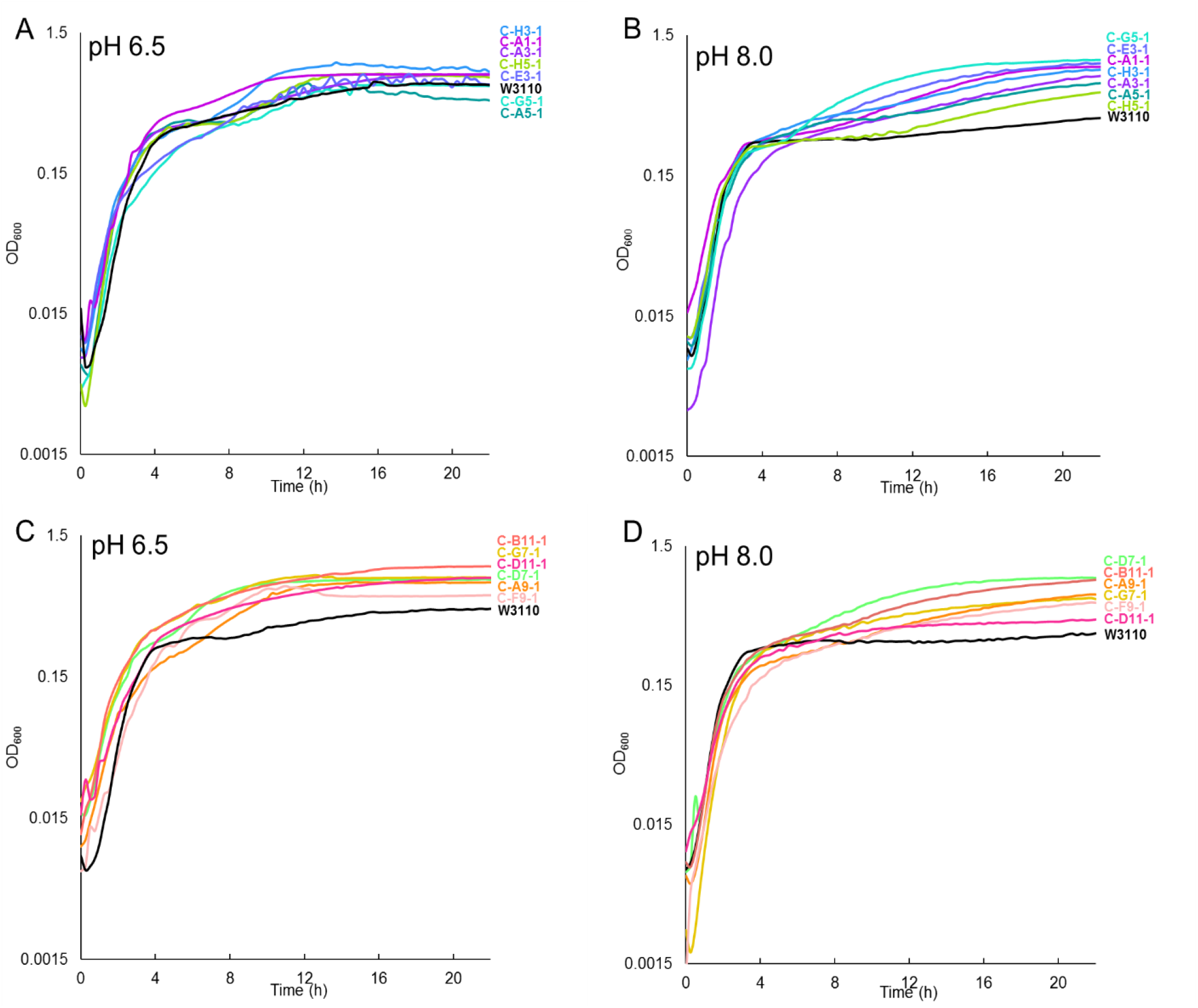
Low pH CCCP-evolved strains cultured over 22 hours at OD600 in (**A**) 100 mM K-PIPES pH 6.5 and (**B**) 100 mM TAPS pH 8.0. In 100 mM TAPS pH 8.0, W3110 differs significantly from all strains but H5-1 at p <0.05 (ANOVA with Tukey post hoc, F=34.698, p < 0.001, n=8 for each strain). High pH CCCP-evolved strains were tested over 22 hours at OD600 in (**C**) 100 mM K-PIPES pH 6.5 and (**D**) 100 mM TAPS pH 8.0. Each curve represents the median 16-h culture density of eight replicates.

In order to test whether these improvements in growth reflect some other factor associated with the stress of growing in a microplate, we conducted an aerated growth curve in baffled flasks. We selected two evolved strains from the low pH and high pH conditions and measured their grown compared to the ancestor at pH 6.5 100 mM PIPES and pH 8.0 100 mM TAPS, respectively, over the course of 8 hours using a traditional growth assay conducted in comparatively more aerobic baffled flasks. We saw no significant growth differences between ancestral and evolved populations (data not shown).

To determine whether conferred growth advantages to pH stress would occur under more extreme acidic and basic conditions, growth of CCCP-evolved strains was tested at pH 5.0 100 mM MES (**Fig. 8A**) and pH 9.0 100 mM AMPSO (**Fig. 8B**). Tests at pH 5.0 showed no significant differences. A pH 9.0, the pH 8.0-evolved strains (C-B11-1, C-D11-1, C-F9-1, C-G7-1) grew to a higher density than the ancestor while the pH 6.5-evolved strains (C-A1-1, C-A3-1, C-G5-1) grew less well than the ancestor (**Fig. 8B**). These results show some shifting of growth range in favor of the pH during serial culture. At pH 7.0, CCCP-evolved strains showed marginal differences from the ancestor (**Fig. 9**).

**Figure 8.**
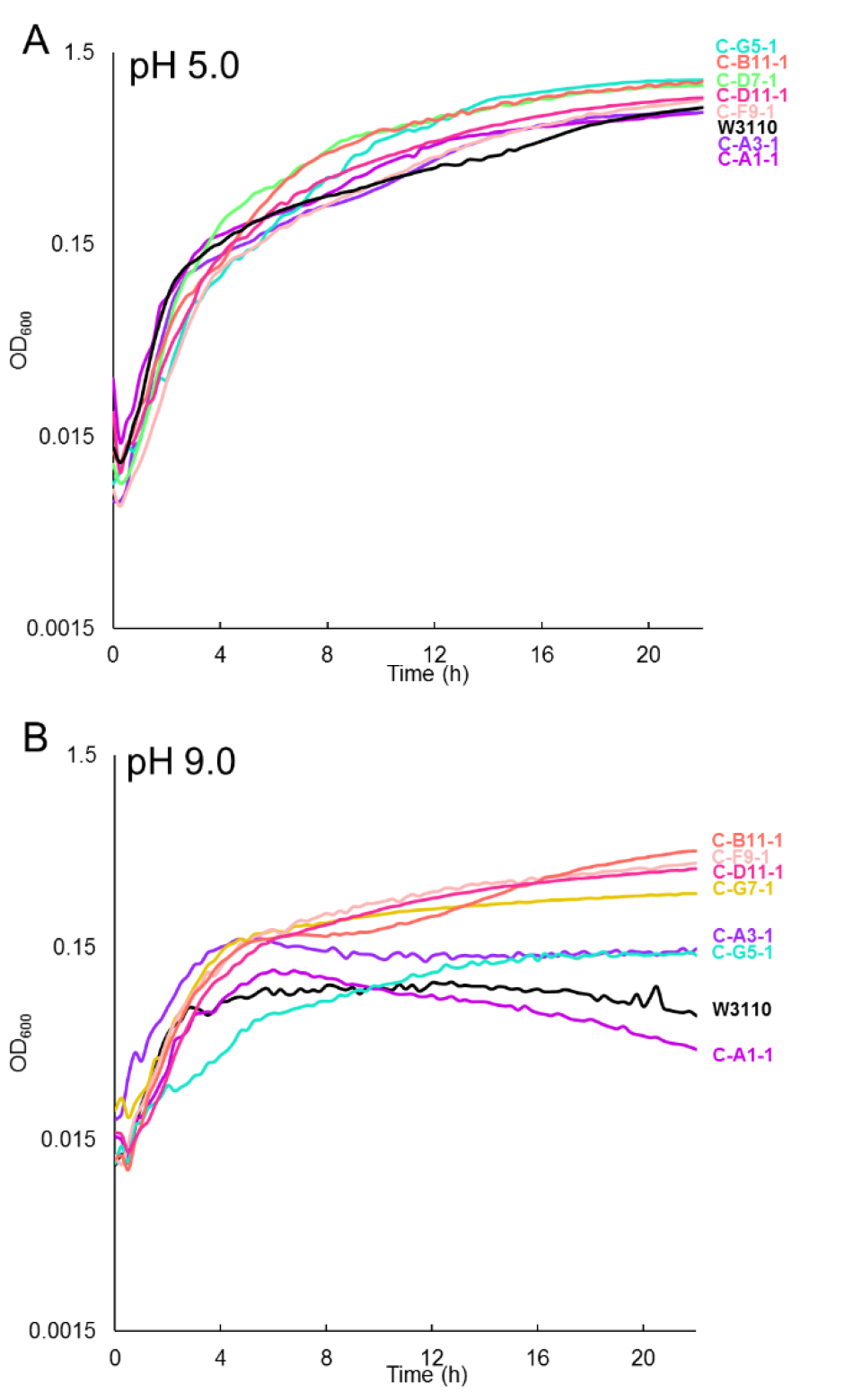
CCCP-evolved strains tested over 22 hours at OD_600_ in (**A**) 100 mM MES pH 5.0 and (**B**) 100 mM AMPSO pH 9.0. In 100 mM MES pH 5.0, W3110 differs significantly from all evolved strains except A3-1 at p < 0.05 (ANOVA with Tukey post hoc, F=23.481, p < 0.001, n=8 for each strain). In 100 mM AMPSO pH 9.0, all low pH CCCP-evolved strains differ from all high pH strains at p <0.05 (ANOVA with Tukey post hoc, F=34.9, p=2e^-16^, n=8 for each strain).

**Figure 9.**
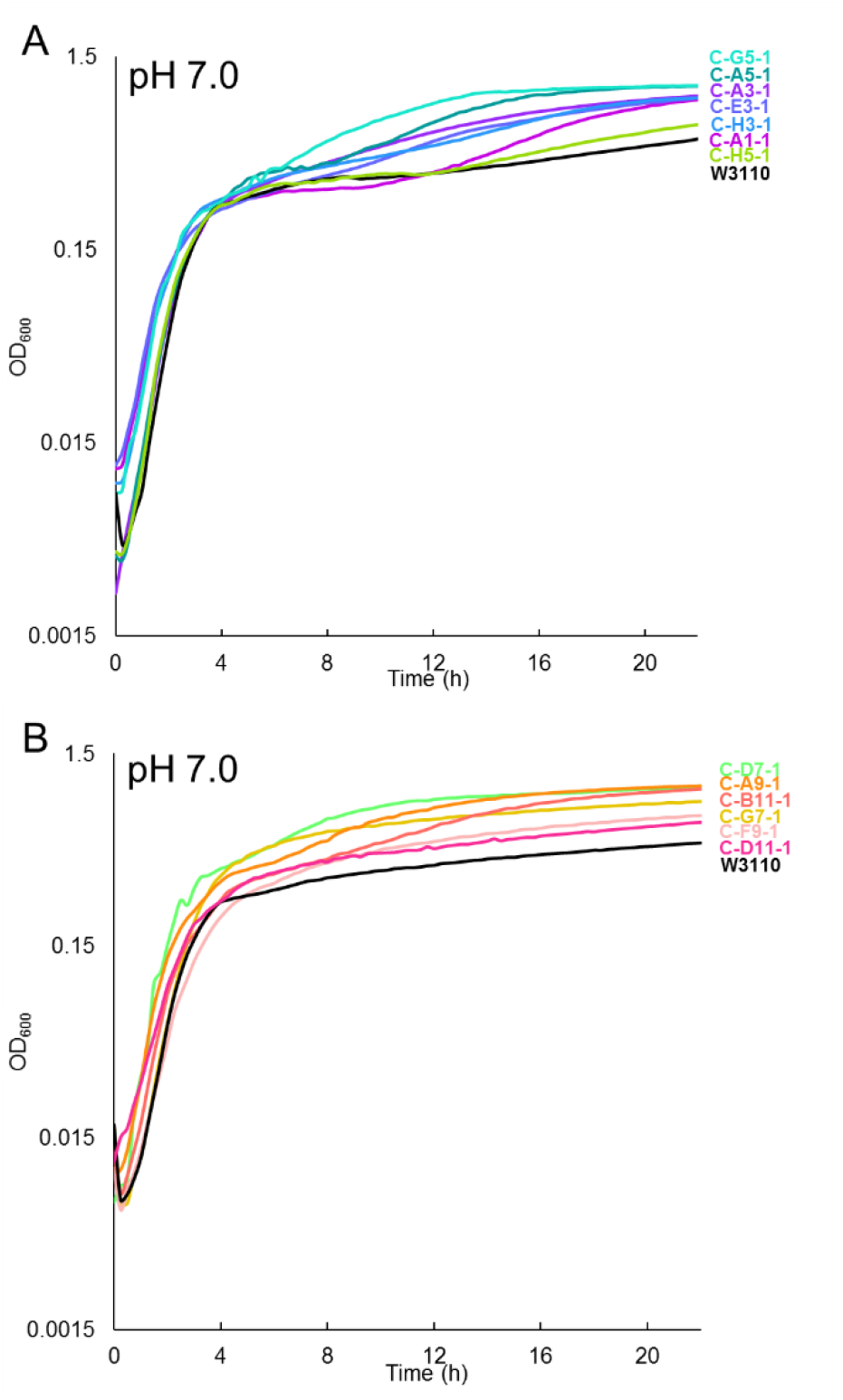
(**A**) Low pH CCCP-evolved strains in pH 7.0 100 mM MOPS over 22 hours at OD600. W3110 differs significantly from all strains but H5-1 at p<0.05 (ANOVA with Tukey post hoc, F=15.925, p < 0.001, n=8 for each strain). (**B**) High pH CCCP-evolved strains in pH 7.0 100 mM MOPS over 22 hours at OD600. W3110 differs significantly from A9-1 and B11-1 at p <0.05 (ANOVA with Tukey post hoc, F=10.721, p < 0.001, n=8 for each strain).

### Evolved strains show many mutations to *adhE*

Multiple mutations were found in *adhE* (acetaldehyde-CoA dehydrogenase) in the strains that evolved in CCCP at pH 8.0. The CCCP stock had been dissolved in ethanol, leading to 1.5% ethanol at the highest concentration used in growth media (150 μM CCCP for both pH 8.0 and pH 6.5). We tested the ability of the high pH evolved strains to grow in 1.5% ethanol, and found that evolved strains grew to a significantly higher optical density than the ancestor (**Fig. 10A**). We also tested the CCCP tolerance of our strains using a stock dissolved in DMSO, instead of ethanol. The CCCP fitness advantage remained in the absence of ethanol (**Fig. 10B**).

**Figure 10.**
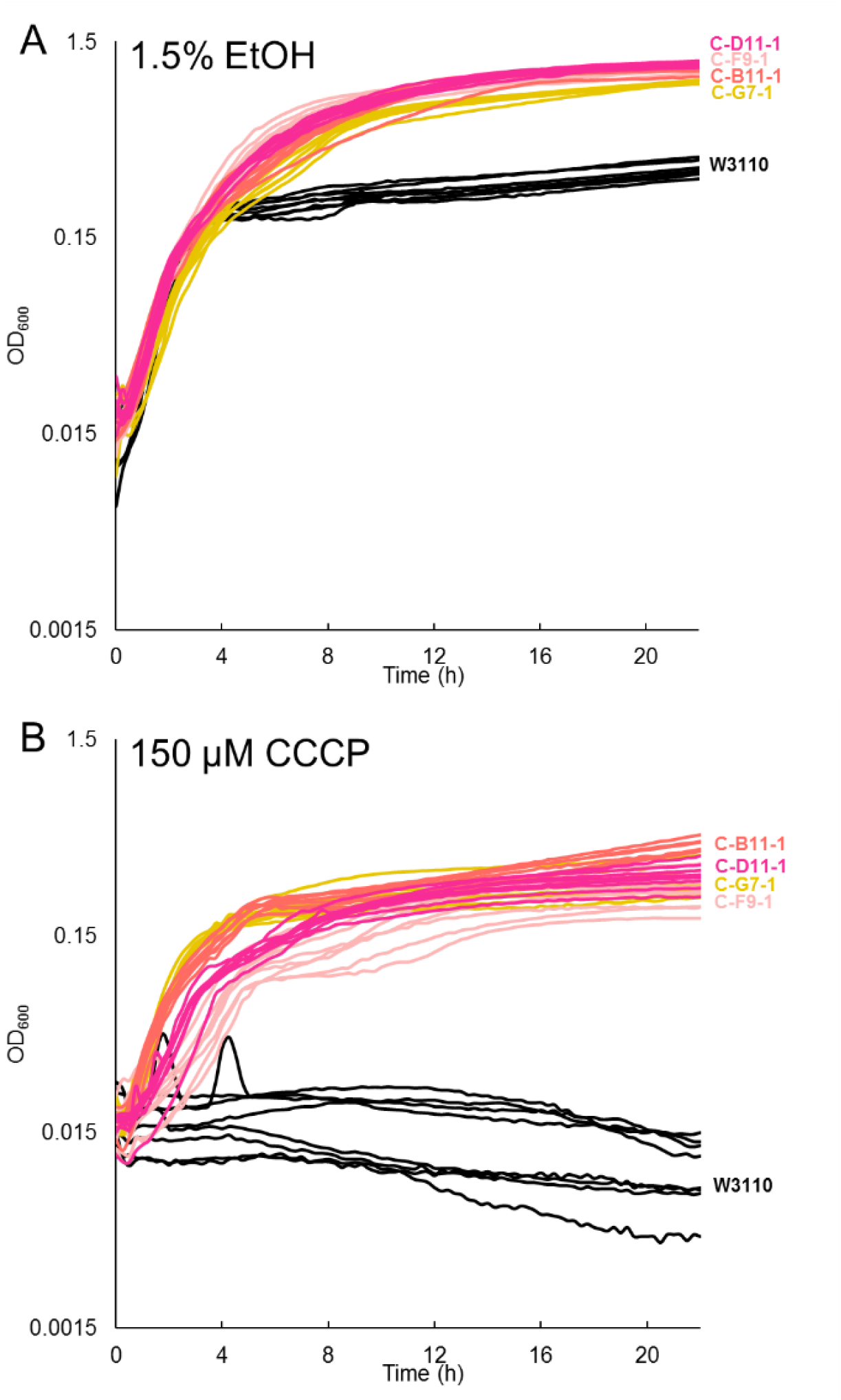
Growth of high pH CCCP-evolved strains in LBK 100 mM TAPS pH 8.0 with (**A**) 1.5% ethanol or (**B**) 150 μM CCCP dissolved in DMSO.

## DISCUSSION

A major finding of our CCCP experimental evolution of *E. coli* was that all strains but one (C-B11-1) showed mutations to *mprA,* the repressor of *emrAB.* The *emrAB* genes also showed point mutations in many of the strains. The *emrAB* genes encode a multidrug efflux transport system that has been previously linked to CCCP resistance (13). Deleting the repressor may increase expression or activity of CCCP efflux via EmrAB-TolC (13, 15). The upregulation of EmrAB in our strains is consistent with their increased resistance to nalidixic acid (**Fig. 6A**), another substrate of the efflux pump (13, 27).

Nevertheless, the strain C-B11-1 that lacked mutations in *emrA* or *mprA* attained comparable fitness increase without increasing CCCP export. This strain showed downregulation of other MDR pumps, via mutations in regulators *gadE* (activates *mtdEF*) or *cecR* (represses *ybhG*; resistance to chloramphenicol and cefoperazone). Other strains showed MDR mutations in MDR components *acrB* and *ybhR.* **Table 4** compiles all the MDR-related mutations we have found to be selected during evolution in CCCP or in benzoate (9). By contrast, microplate experimental evolution under other conditions does not incur loss of multiple drug pumps (10, 11). The one exception would be mutations in the Gad acid fitness island during evolution in near-extreme acid; but acid response components of Gad are the likely targets.

**Table 4.**
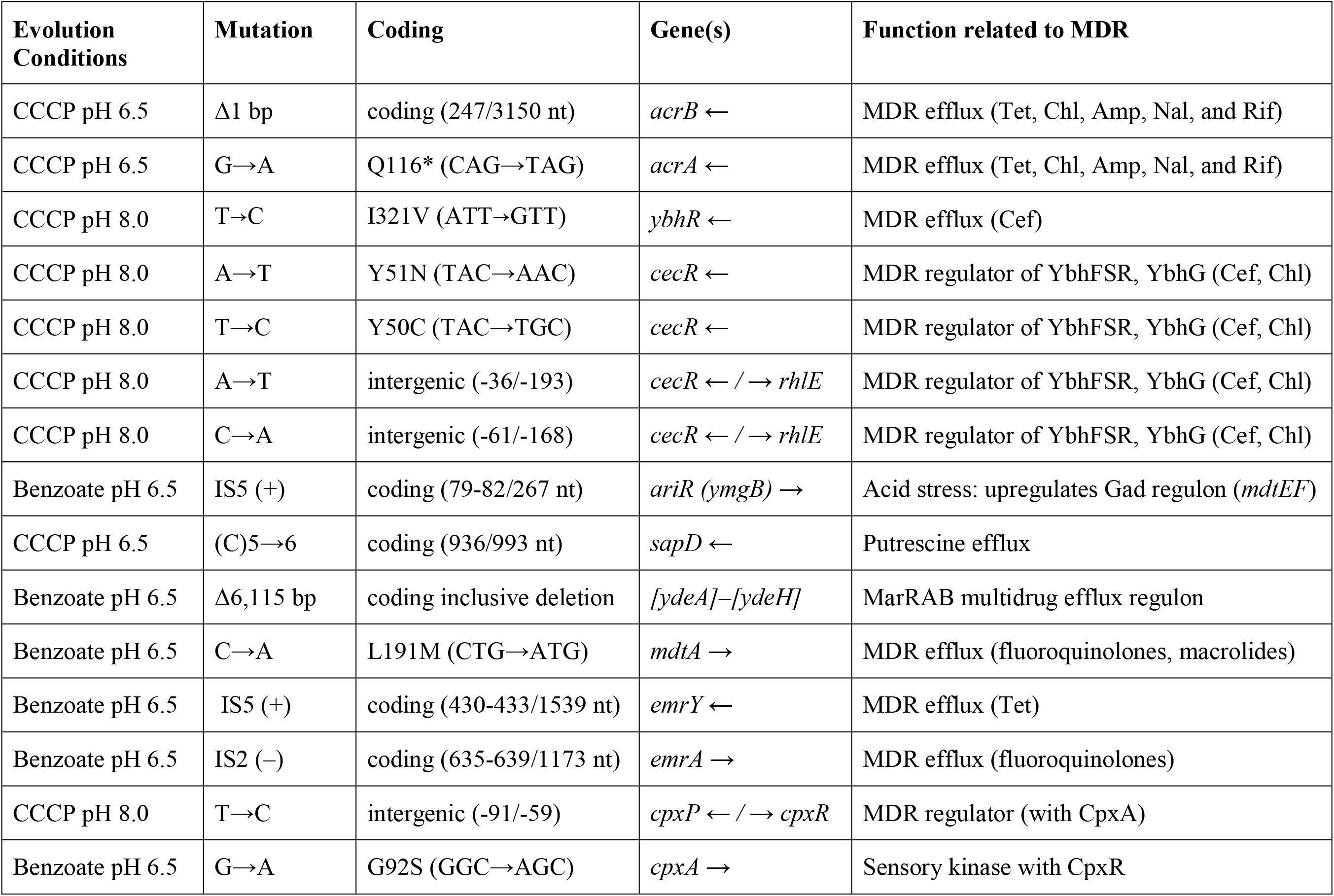
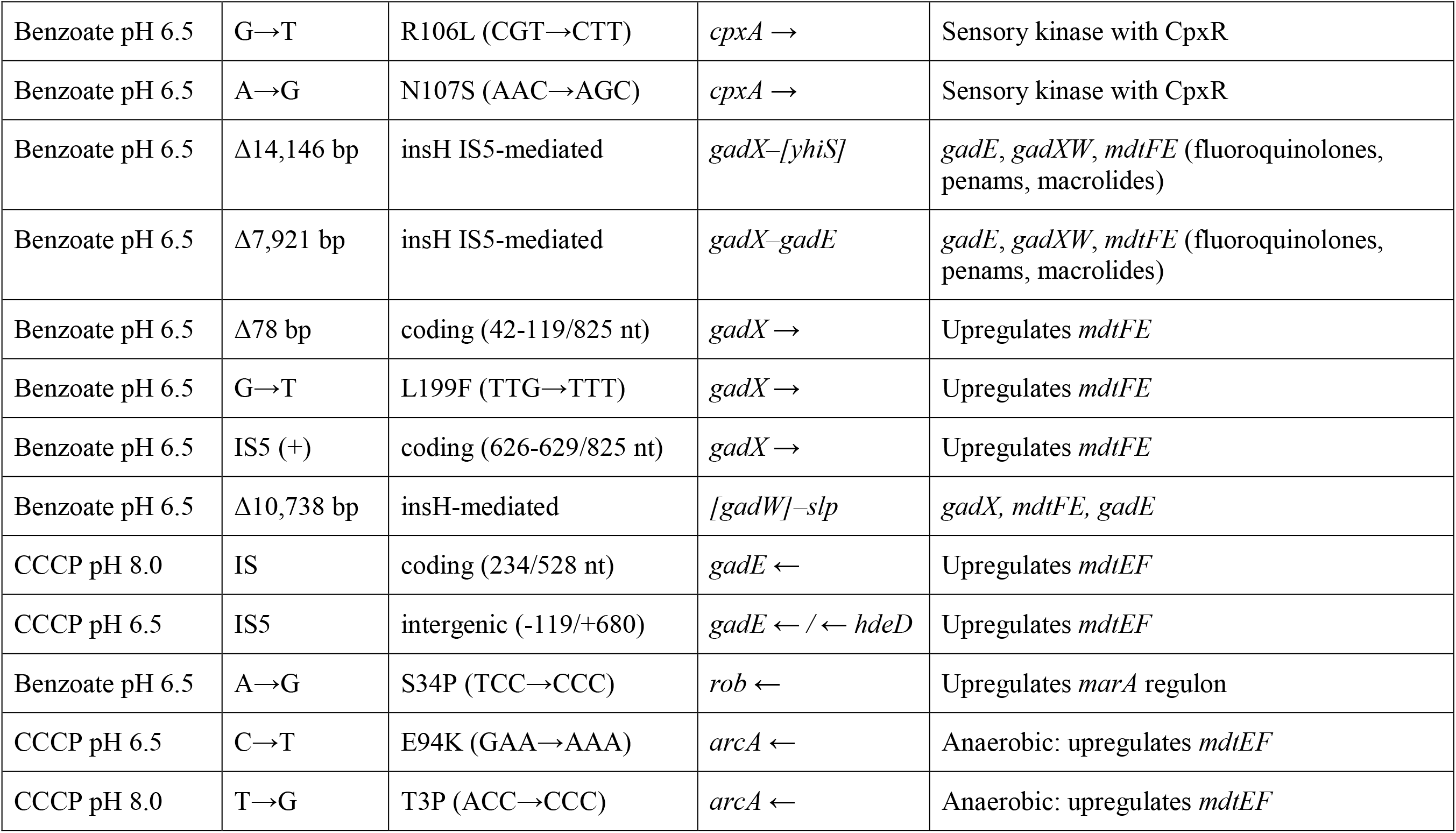
Multi-drug pumps and regulators showing mutations selected by long-term exposure to benzoate or to CCCP.*

Overall, our results add to a pattern of selection against drug efflux pumps and their regulators under evolution with an uncoupler or a partial uncoupler (benzoate). However, unlike CCCP, benzoate acts at relatively high aqueous concentrations that equilibrate across the membrane. These concentrations cannot be effluxed by MDR pumps; therefore, adaptation to the presence of benzoate and other permeant acids may be more dependent upon mutations that decrease transmembrane proton flux through unneeded pumps. We are following up this work with more extended comparison of the relative fitness costs of MDR genes, using flow cytometry competition assays (28).

## MATERIALS AND METHODS

### CCCP Experimental Evolution a pH 6.5 and 8.0

For experimental evolution, *E. coli* K-12 W3110 populations were cultured at 37°C in a 96-well microplate in buffered LBK media (8). To compare the effects of CCCP at low versus high pH, 24 populations with 20 μM CCCP were buffered at pH 6.5 with 100 mM Na-PIPES (which contained 0.55 M PIPES buffer and 1.4 M NaOH), and 24 populations with 50 μM CCCP were buffered at pH 8.0 with 100 mM TAPS. The populations grew to stationary phase, and after a total of 22 h were diluted 1:100 daily. A multi-channel pipettor with filter tips was used for daily transfer of 2-μl culture into 200-μl fresh medium. Over the course of their serial dilutions, the populations’ concentration of CCCP was increased, with the pH 6.5 condition increasing from 20 μM to 150 μM and the pH 8.0 condition increasing from 50 μM to 150 μM CCCP (**Table 1**).

After the populations had grown for 1,000 generations, clonal isolates were obtained from selected well populations. Of the populations evolved at pH 6.5, seven isolates from different wells were selected for full genome sequencing and of those at pH 8.0, six isolates were selected for whole genome sequencing. Clonal isolates were generated by streaking samples from all wells three times on LBK plates. Each isolate was given a population designation: “C” for CCCP evolution, the alphanumeric position in the well plate, and a digit for the isolate number from the given well population.

Whole Genome Sequencing and Sequence Analysis. The genomes of fourteen isolates from generation 1,000 and the ancestor were sequenced (**Table S1**). DNA extraction was performed using the Epicentre Masterpure Purification Kit. The genomes were validated and quantitated using the Illumina TruSeq Nano DNA Library Preparation Kit by the Research and Technology Support Facility at Michigan State University. Sequencing was then performed utilizing the Illumina MiSeq platform in 2x300 bp format, resulting in 20-25 million read pairs. Read alignment and mutation predictions were performed using the computational pipeline *breseq* (v.0.28.1, v.0.30.0, and v.0.30.2) with default parameters. The reads were mapped to the *E. coli* K-12 W3110 reference (NC_007779.1). Mutations found in both the ancestor and evolved strain were ignored.

### Knockouts in genes of interest

To determine the fitness impact of specific genes on growth in CCCP, certain genes were deleted from both the ancestor and evolved strains. Donor *E. coli* strains containing *kanR* insertion knockouts were obtained from the Keio collection (29). Deletion strains of W3110 background were constructed by P1 phage transduction (9). Knockout constructs were confirmed by PCR amplification of the *kanR* insert and were checked against the recipient and donor strains using X-Gal for *lacZ*^+^ (W3110 background). Colony PCR was performed using Lucigen CloneID Colony PCR Master Mix to determine whether the *kanR* cassette had been inserted into the correct gene and that the recipient strain was as expected. Primers included gene of interest forward direction, gene of interest reverse direction, internal KAN gene forward, internal KAN gene reverse, and Primer KT for within the KAN marker.

### Recombineering

In order to generate strain C-B11-1 *rng*^+^, a section of gene *rng* containing the non-sense mutation in CCCP-evolved isolate C-B11-1 was replaced with the wild-type sequence using recombineering (30, 31). A counter-selectable *cat-sacB* cassette was amplified from colonies of *E. coli* strain XLT241 using primers with 5’ homology to regions surrounding the *rng* mutation site in C-B11-1 (5’-CAGCGCCGAAAAACCATTAACGCTGGTTTTCACCCGGTCTTGTGACGGAAGATCACTTCGCAGAATA-3’ and 5’-ATAATGAAGATCACCGCCGCCGAGTGCTGCACTCGCTGGAGCAATCAAAGGGAAAACTGTCCATAT-3’). Isolate C-B11-1 transformed with the heat-inducible pSIM6∷ampR plasmid was cultured to mid-log phase at 32°C then transferred to 42°C to induce recombineering proteins. Cells were then made electrocompetent and electroporated with the *cat-sacB* PCR product containing edges homologous to regions flanking the *rng* mutation site in B11-1. The cells were outgrown at 32°C for 3 to 5 hours and plated on LB with 10 μg/ml chloramphenicol. Colonies were then screened for chloramphenicol resistance and sucrose sensitivity. To replace the *cat-sacB* cassette with the wild-type *rng* sequence, successful colonies were made electrocompetent and electroporated with a DNA oligonucleotide containing the wild-type sequence of the region replaced by *cat-sacB* as well as homology to the flanking regions (5’-CCATTAACGCTGGTTTTCACCCGGTCTTTGCTCAACGCCTGCTCCAGCGAGTGCAGCACTCGGCGGCGGTG-3’). The cells were then plated on LB lacking NaCl and containing 6% sucrose. Colonies were further screened for resistance to sucrose and chloramphenicol sensitivity. The *rng* sequence of the new construct was then confirmed by PCR sequencing (5’-GCATGGTGGACACGAACAAT-3’ and 5’-CGCTGGAACGCAAAGTAGAA-3’).

### Growth Assays

Batch culture growth was assayed under semiaerobic conditions based on previous procedures (9–11). A 96-well microplate was filled with the desired media and inoculated at a concentration of 1:200 from an overnight culture in each test. Eight replicates from two independent overnight cultures were performed for each strain within a single plate, and growth curves were then run in triplicate. Microplates were placed in the Spectramax 384 spectrophotometer and cultured for 22 hours at 37°C with OD_600_ measured every 15 min.

Growth curves for antibiotic tests were inoculated from overnight cultures that were grown in LBK 100 mM MOPS pH 7.0. These antibiotic growth curves were tested in LBK 100 mM MOPS pH 7.0 with 1 μg/ml ampicillin, 1 μg/ml tetracycline, 4 μg/ml chloramphenicol or 6 μg/ml nalidixic acid.

For growth curves under aeration, 125-ml baffled flasks containing 20 ml of LBK (pH 8.0 100 mM TAPS or pH 6.5 100 mM Na-PIPES) were inoculated 1:200 from overnight cultures. The cultures were grown in a water bath shaker set to 37 °C 200 rpm. 200-μl aliquots were removed from the baffled flasks and read in a 96 well plate (wavelength 600 nm) every thirty minutes for eight hours in order to construct the growth curve.

### Statistics

Means are reported with standard error of the mean (SEM). Statistics were computed using R packages. Samples were compared using Analysis of Variance tests (ANOVA) with Tukey post hoc tests.

### Accession number

Sequence data have been uploaded in the NCBI Sequence Read Archive (SRA) under accession number SRP157768.

## ACKNOWLEDGMENTS

This work was supported by award MCB-1613278 from the National Science Foundation and the support of Kenyon College. We thank Ellen Broeren for excellent technical support.

## REFERENCES

1. Mitchell P. 2011. Chemiosmotic coupling in oxidative and photosynthetic phosphorylation. Biochim Biophys Acta-Bioenerg 1807:1507–1538.

2. Kasianowicz J, Benz R, McLaughlin S. 1984. The kinetic mechanism by which CCCP (carbonyl cyanide m-Chlorophenylhydrazone) transports protons across membranes. J Membr Biol 82:179–190.

3. McLaughlin SG, Dilger JP. 1980. Transport of protons across membranes by weak acids. Physiol Rev 60:825–863.

4. Gould JM. 1979. Respiration-linked proton transport, changes in external pH, and membrane energization in cells of *Escherichia coli*. J Bacteriol 138:176–184.

5. Lobritz MA, Belenky P, Porter CBM, Gutierrez A, Yang JH, Schwarz EG, Dwyer DJ, Khalil AS, Collins JJ. 2015. Antibiotic efficacy is linked to bacterial cellular respiration. Proc Natl Acad Sci 112:8173–8180.

6. Diez-Gonzalez F, Russell JB. 1997. Effects of carbonylcyanide-m-chlorophenylhydrazone (CCCP) and acetate on *Escherichia coli* O157:H7 and K-12: Uncoupling versus anion accumulation. FEMS Microbiol Lett 151:71–76.

7. MacLeod RA, Wisse GA, Stejskal FL. 1988. Sensitivity of some marine bacteria, a moderate halophile, and *Escherichia coli* to uncouplers at alkaline pH. J Bacteriol 170:4330–4337.

8. Harden MM, He A, Creamer K, Clark MW, Hamdallah I, Martinez KA, Kresslein RL, Bush SP, Slonczewski JL. 2015. Acid-adapted strains of *Escherichia coli* K-12 obtained by experimental evolution. Appl Environ Microbiol 81:1932–1941.

9. Creamer KE, Ditmars FS, Basting PJ, Kunka KS, Hamdallah IN, Bush SP, Scott Z, He A, Penix SR, Gonzales AS, Eder EK, Camperchioli DW, Berndt A, Clark MW, Rouhier KA, Slonczewski JL. 2017. Benzoate-and salicylate-tolerant strains of *Escherichia coli* K-12 lose antibiotic resistance during laboratory evolution. Appl Environ Microbiol 83:e02736.

10. He A, Penix SR, Basting PJ, Griffith JM, Creamer KE, Camperchioli D, Clark MW, Gonzales AS, Sebastian Chávez Erazo J, George NS, Bhagwat AA, Slonczewski JL. 2017. Acid evolution of *Escherichia coli* K-12 eliminates amino acid decarboxylases and reregulates catabolism. Appl Environ Microbiol 83:e00442.

11. Hamdallah I, Torok N, Bischof KM, Majdalani N, Chadalavada S, Mdluli N, Creamer KE, Clark M, Holdener C, Basting PJ, Gottesman S, Slonczewski JL. 2018. Experimental evolution of *Escherichia coli* K-12 at high pH and RpoS induction. Appl Environ Microbiol 84:e00520.

12. Deng Z, Shan Y, Pan Q, Gao X, Yan A. 2013. Anaerobic expression of the *gadE-mdtEF* multidrug efflux operon is primarily regulated by the two-component system ArcBA through antagonizing the H-NS mediated repression. Front Microbiol 4:194.

13. Lomovskaya O, Lewis K, Matin A. 1995. EmrR is a negative regulator of the *Escherichia coli* multidrug resistance pump EmrAB. J Bacteriol 177:2328–2334.

14. Lomovskaya O, Lewis K. 1992. *emr*,an *Escherichia coli* locus for multidrug resistance. Proc Natl Acad Sci U S A 89:8938–8942.

15. Nikaido H. 1996. Multidrug efflux pumps of gram-negative bacteria. J Bacteriol 178:5853–5859.

16. Xiong A, Gottman A, Park C, Baetens M, Pandza S, Matin A. 2000. The EmrR protein represses the *Escherichia coli emrRAB* multidrug resistance operon by directly binding to its promoter region. Antimicrob Agents Chemother 44:2905–2907.

17. Brooun A, Tomashek JJ, Lewis K. 1999. Purification and ligand binding of EmrR, a regulator of a multidrug transporter. J Bacteriol 181:5131–5133.

18. Gage DJ, Neidhardt FC. 1993. Adaptation of *Escherichia coli* to the uncoupler of oxidative phosphorylation 2,4-dinitrophenol. J Bacteriol 175:7105–7108.

19. Yamanaka Y, Shimada T, Yamamoto K, Ishihama A. 2016. Transcription factor CecR (YbiH) regulates a set of genes affecting the sensitivity of *Escherichia coli* against cefoperazone and chloramphenicol. Microbiology 162:1253–1264.

20. Hommais F, Krin E, Coppée JY, Lacroix C, Yeramian E, Danchin A, Bertin P. 2004. GadE (YhiE): A novel activator involved in the response to acid environment in *Escherichia coli*. Microbiology 150:61–72.

21. Seo SW, Kim D, O”Brien EJ, Szubin R, Palsson BO. 2015. Decoding genome-wide GadEWX-transcriptional regulatory networks reveals multifaceted cellular responses to acid stress in *Escherichia coli*. Nat Commun 6:7970.

22. Zhang Y, Xiao M, Horiyama T, Zhang Y, Li X, Nishino K, Yan A. 2011. The multidrug efflux pump MdtEF protects against nitrosative damage during the anaerobic respiration in *Escherichia coli*. J Biol Chem 286:26576–26584.

23. Roggiani M, Yadavalli SS, Goulian M. 2017. Natural variation of a sensor kinase controlling a conserved stress response pathway in *Escherichia coli*. PLoS Genet 13:1007101.

24. Li X, Gu Y, Dong H, Wang W, Dong C. 2015. Trapped lipopolysaccharide and LptD intermediates reveal lipopolysaccharide translocation steps across the *Escherichia coli* outer membrane. Sci Rep 5:11883.

25. Okusu H, Ma D, Nikaido H. 1996. AcrAB efflux pump plays a major role in the antibiotic resistance phenotype of *Escherichia coli* multiple-antibiotic-resistance (Mar) mutants. J Bacteriol 178:306–308.

26. Baucheron S, Tyler S, Boyd D, Mulvey MR, Chaslus-dancla E, Cloeckaert A. 2004. AcrAB-TolC directs efflux-mediated multidrug resistance in *Salmonella enterica* serovar Typhimurium DT104. Antimicrob Agents Chemother 48:3729–3735.

27. Sulavik MC, Houseweart C, Cramer C, Jiwani N, Murgolo N, Greene J, DiDomenico B, Shaw KJ, Miller GH, Hare R, Shimer G. 2001. Antibiotic susceptibility profiles of *Escherichia coli* strains lacking multidrug efflux pump genes. Antimicrob Agents Chemother 45:1126–1136.

28. Gullberg E, Cao S, Berg OG, Ilbäck C, Sandegren L, Hughes D, Andersson DI. 2011. Selection of resistant bacteria at very low antibiotic concentrations. PLoS Pathog 7:e1002158.

29. Baba T, Ara T, Hasegawa M, Takai Y, Okumura Y, Baba M, Datsenko KA, Tomita M, Wanner BL, Mori H. 2006. Construction of *Escherichia coli* K-12 in-frame, single-gene knockout mutants: the Keio collection. Mol Syst Biol 2:8.

30. Sharan SK, Thomason LC, Kuznetsov SG, Court DL. 2009. Recombineering: a homologous recombination-based method of genetic engineering. Nat Protoc 4:206–223.

31. Thomason LC, Sawitzke JA, Li X, Costantino N, Court DL. 2014. Recombineering: genetic engineering in bacteria using homologous recombination, p.1.16.1-1.16.39. In Current Protocols in Molecular Biology.

